# Error-prone bypass of DNA lesions during lagging strand replication is a common source of germline and cancer mutations

**DOI:** 10.1101/200691

**Authors:** Vladimir B. Seplyarskiy, Evgeny E. Akkuratov, Natalia V. Akkuratova, Maria A. Andrianova, Sergey I. Nikolaev, Georgii A. Bazykin, Igor Adameyko, Shamil R. Sunyaev

## Abstract

Spontaneously occurring mutations are of great relevance in diverse fields including biochemistry, oncology, evolutionary biology, and human genetics. Studies in experimental systems have identified a multitude of mutational mechanisms including DNA replication infidelity as well as many forms of DNA damage followed by inefficient repair or replicative bypass. However, the relative contributions of these mechanisms to human germline mutations remain completely unknown. Here, based on the mutational asymmetry with respect to the direction of replication and transcription, we suggest that error-prone damage bypass on the lagging strand plays a major role in human mutagenesis. Asymmetry with respect to transcription is believed to be mediated by the action of transcription-coupled DNA repair (TC-NER). TC-NER selectively repairs DNA lesions on the transcribed strand; as a result, lesions on the non-transcribed strand are preferentially converted into mutations. In human polymorphism we detect a striking similarity between transcriptional asymmetry and asymmetry with respect to replication fork direction. This parallels the observation that damage-induced mutations in human cancers accumulate asymmetrically with respect to the direction of replication, suggesting that DNA lesions are asymmetrically resolved during replication. Re-analysis of XR-seq data, Damage-seq data and cancers with defective NER corroborate the preferential error-prone bypass of DNA lesions on the lagging strand. We experimentally demonstrate that replication delay greatly attenuates the mutagenic effect of UV-irradiation, in line with the key role of replication in conversion of DNA damage to mutations. We conservatively estimate that at least 10% of human germline mutations arise due to DNA damage rather than replication infidelity. The number of these damage-induced mutations is expected to scale with the number of replications and, consequently, paternal age.

---

Experiments in well-controlled genetic systems and in model organisms have uncovered that DNA polymerases make errors and resulting mismatches become mutations^1^. An alternative mechanism of mutagenesis due to misrepaired DNA damage or DNA damage bypassed by translesion (TLS) polymerases has been extensively studied in experimental systems exposed to exogenous mutagens^2,3^. Although these studies do shed light on the mechanistic details of mutagenesis, well-controlled experimental systems provide little information on the relative contributions of these mechanisms to naturally occurring human mutations. More recently, computational genomics approaches have revealed statistical properties of mutations occurring in the germline^4–7^, in tumors^8^ and in embryo during early stages of development^9–11^. In cancer, many types of mutations have been successfully attributed to the action of specific mutagenic forces^12^. A number of studies have explored how cancer mutations scale with age at diagnosis^13,14^ and how human germline mutations scale with paternal age^15–17^. It was hypothesized that the dependency of the number of accumulated mutations on the number of cell divisions may also reflect the replicative origin of mutations^18,19^. However, a quantitative model suggests that accumulation of both damage-induced and co-replicative mutations may scale with the number of cell divisions^20^. Therefore, we still do not know whether DNA damage substantially contributes to heritable human mutations or whether natural mutagenesis in humans is mostly due to errors in replication.

To discriminate between co-replicative mutations and damage-induced mutations, we rely on statistical properties of mutations unequivocally associated with DNA damage. Both germline and cancer mutations leave footprints in the form of mutational asymmetry with respect to the direction of transcription (T-asymmetry). T-asymmetry reflects the prevalence of mutations that originate from lesions on the non-transcribed strand that could not be repaired by TC-NER^21,22^. Thus, the analysis of T-asymmetry may be used to quantify the prevalence of mutations arising from DNA lesions. Genomic data on cancers in which most mutations are caused by the action of specific, well-understood, DNA-damage-inducing agents provide an additional perspective on properties of damage-induced mutations. Notably, the level of T-asymmetry in these cancers is exceptionally high.

The most obvious statistical feature associated with replication is asymmetry with respect to the direction of the replication fork (R-asymmetry). R-asymmetry may reflect differential fidelity of replication between the leading and lagging strands. Alternatively, R-asymmetry may be caused by the strand-specific bypass of DNA damage. Bulky DNA lesions not repaired prior to replication can either lead to fork regression followed by error-free repair or be bypassed by TLS polymerases^23,24^. TLS synthesis is error-prone. It does not remove the lesion and commonly introduces mutations on the newly synthesized strand. It has been asserted that the error-prone bypass process has different properties on leading and lagging strands^23,25^ that would lead to R-asymmetry.

As a starting point, we compare R-asymmetry with T-asymmetry. To avoid the interference of statistical signals between the two types of asymmetries, R-asymmetry is estimated exclusively in intergenic regions and T-asymmetry only for genic regions. We calculate R-asymmetries for the 92 types of single-nucleotide mutations (excluding NpCpG>NpTpG mutations) in each trinucleotide context. C>T mutations in the NpCpG context are excluded because they usually arise via conversion of methylated cytosines directly into thymines by deamination^26^ (Supplementary Note 1). Figure 1 shows data for rare (allele frequency below 0.1%) SNVs from the gnomAD dataset. Supplementary Figure 1 shows that R-asymmetries across different contexts are concordant between rare SNPs and *de novo* mutations.

**Figure 1.**
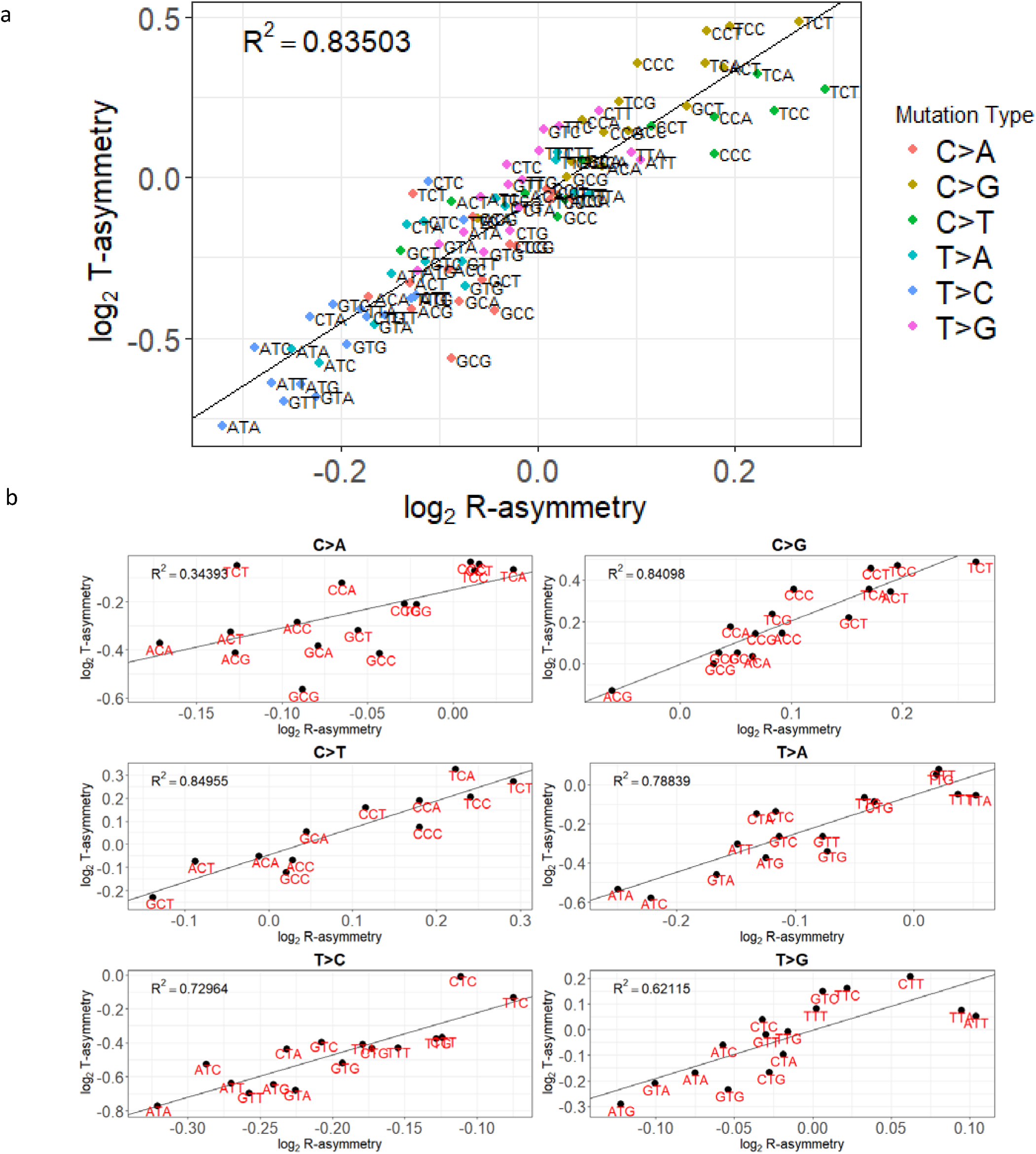
R-asymmetry and T-asymmetry patterns in human polymorphism. **a**, Relationship between R-asymmetry and T-asymmetry for 92 mutation types (NpCpG>T mutations excluded). b, Relationship between R-asymmetry and T-asymmetry shown separately for the six types of single-nucleotide mutations to highlight the effects of adjacent nucleotides.

Strikingly, there is a high concordance between T-asymmetry and R-asymmetry across mutation types in both tri-nucleotide contexts (Figure 1) and penta-nucleotide contexts (Supplementary Figure 2). Mutation types that are predominant on the lagging strand are also more common on the non-transcribed strand (Figure 1a; R^2^=0.84; p-value=5.6*10^−37^). Moreover, this association holds even when six basic mutation classes are considered separately (Figure 1b).

As noted above, T-asymmetry arises from DNA damage on the non-transcribed strand that is invisible to TC-NER repair^6^. The unrepaired DNA lesions are occasionally converted into mutations. As a result, mutation types commonly induced by damage are biased towards the non-transcribed strand, and the level of T-asymmetry scales with the proportion of damage-induced mutations in the context. Figure 1 suggests that R-asymmetry may be due to similarly differential resolution of DNA damage between leading and lagging strands. DNA lesions on the lagging strand would be more frequently converted into mutations, probably due to error-prone damage bypass.

To follow up on this hypothesis, we analyze R-asymmetry in cancer genomes that have been influenced by specific mutagens. Four cancer types in PCAWG datasets contain samples with high levels of T-asymmetry in specific mutation contexts: melanoma, predominated by UV-induced C>T mutations (signature 7)^8^; two lung cancers (LUAD and LUSC), predominated by smoking-induced G>T mutations (signature 4); and liver cancer, with a high prevalence of A>G mutations (signatures 12 and 16). All of these processes reflect the action of DNA-damaging mutagens rather than replication infidelity. We find that about 95% of these samples demonstrate a weak but usually significant excess of mutations on the lagging strand during replication (Figure 2, Supplementary Table 1). These three mutagens that damage DNA primarily outside of replication also cause R-asymmetry, strongly suggesting that error-prone bypass on the lagging strand happens frequently. A recent study also found an excess of damage-induced mutations corresponding to COSMIC signatures 23 (unknown etiology) and 24 (Afalotoxin) on the lagging strand^27^. Consistently with our interpretation, a subset of samples lacking mutational signatures associated with DNA damage do not exhibit a lagging strand bias (Supplementary Table 2).

**Figure 2.**
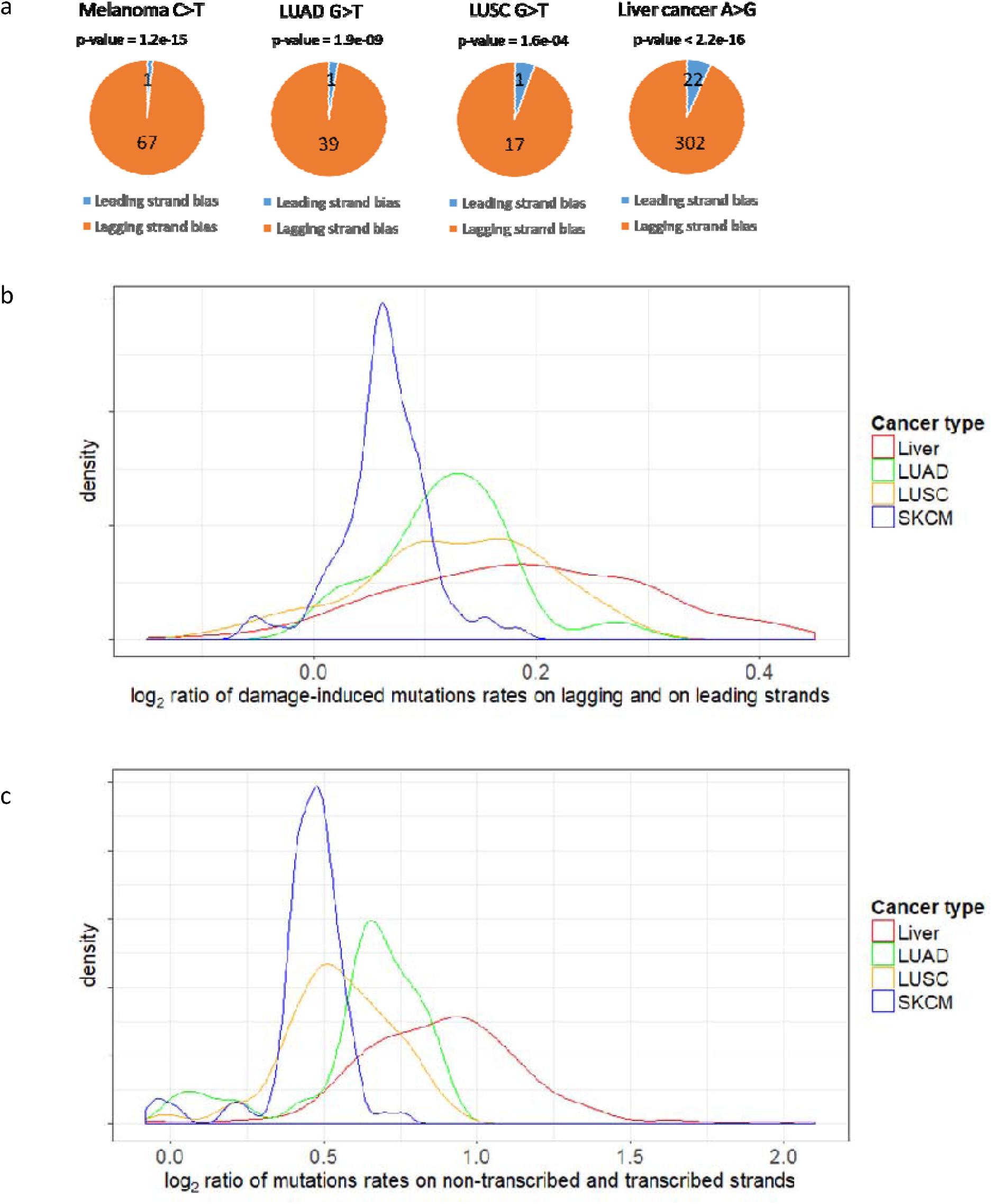
Damage-induced mutations preferentially reside on the lagging strand. **a,** Number of tumor samples among melanomas, lung adeno carcinomas (LUAD), lung squamous carcinomas (LUSC), and liver cancers that have more damage-induced mutations on the leading than on the lagging strand (p-values shown for the goodness-of-fit chi-square test). b,c distribution of R- asymmetry **(b)** and T-asymmetry **(c)** values. Samples with T-asymmetry less than 1.2 were excluded from panel **b**.

The observed R-asymmetry is limited to samples with signatures of bulky damage rather than any type of damage. Tumors with *MUTYH* deficiency have a high load of damage-induced mutations. However, DNA damage in these tumors results in oxo-guanin lesions that do not block progression of RNA and DNA polymerases, and neither T- nor R-asymmetry is detectable in these samples (Supplementary Figure 3). In contrast, R-asymmetry is significantly enhanced in cutaneous squamous cell carcinoma tumors from patients with congenital XPC deficiency (Xeroderma Pigmentosum)^28^. These tumors lack the global genome repair (GG-NER) activity and have elevated levels of bulky damage (See Supplementary Note 2 and Supplementary Figure 4).

If DNA lesions are more frequently bypassed by TLS on the lagging strand directly across the lesion, they will persist on this strand through replication. Therefore, mutational asymmetry caused by the bypass in turn causes the asymmetry of unrepaired DNA damage. We utilize time series XR-seq data^29^ to test whether the activity of the NER system is biased with respect to the replication fork direction. In agreement with the differential bypass hypothesis, repair is more frequently observed on the lagging strand (Figure 3). Moreover, the difference between leading and lagging strands sharply increases with time after UV irradiation as more and more cells complete a round of replication.

**Figure 3.**
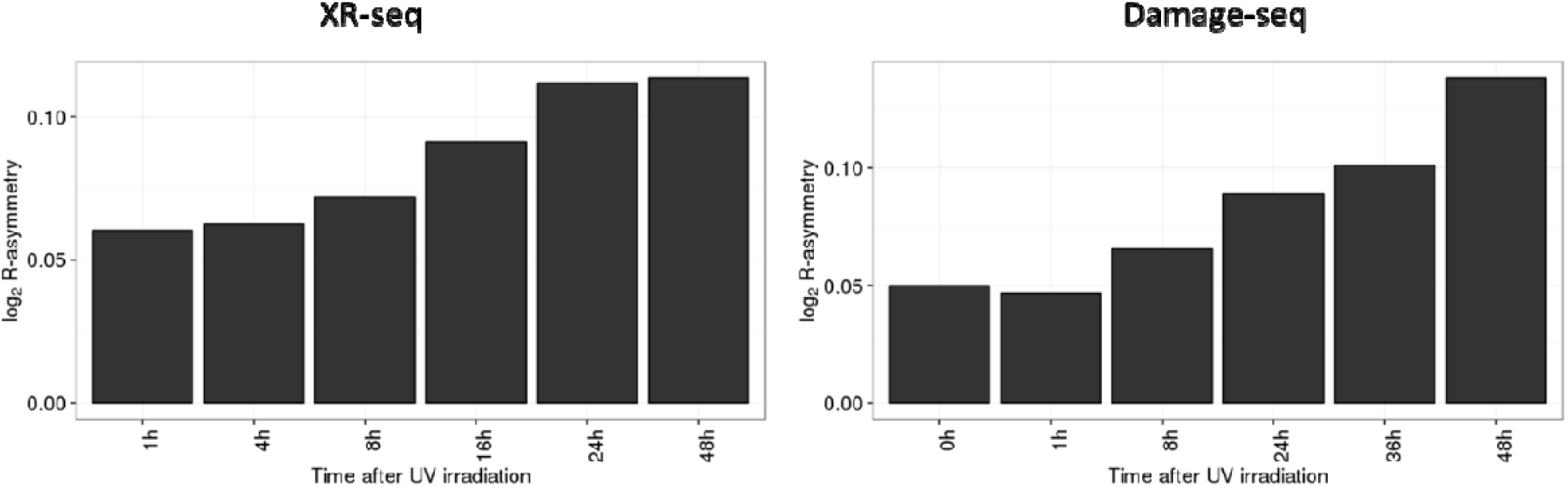
R-asymmetry in UV-irradiated cells. R-asymmetry of repaired CPD damage (left) and CPD damage remaining in DNA (right) as a function of time since UV irradiation.

To test whether the differential activity of the NER system reflects the preferential bypass of DNA damage, we analyze the Damage-seq dataset^30^. Damage-seq detects DNA damage (cyclobutane pyrimidine dimers), and data over a series of time points following the exposure of human fibroblasts to UV radiation are available. The data show a clear dependency on transcription and preferential retention of damage on the non-transcribed strand (Supplementary Figure 5). We observe a lagging strand bias of DNA damage that progressively increases with time, mirroring the trend in XR-seq data (Figure 3).

Collectively, the above observations support the differential replication bypass hypothesis. This hypothesis assumes that many damage-induced mutations do not arise from mis-repair; instead bulky lesions are converted to mutation during DNA replication. Under this assumption, replication delay should reduce mutation rate in cells exposed to damaging agents, because it would provide more time for cells to complete repair. To test this directly, we compared UV-irradiated fibroblast cells exposed and not exposed to roscovitine, which reversibly arrests replication (Figure 4). Colonies that have been grown from fibroblasts not treated with the chemical have ∼14,000 mutations with mutational spectra matching the UV-signature. These UV-induced mutations demonstrated both T- and R-asymmetries quantitatively similar to asymmetries observed in cancer data (*log*_*2*_(T-asym.)=0.50, *log*_*2*_(R-asym.)=0.17, p<0.01 for both). In sharp contrast, colonies derived from UV-irradiated cells that experienced replication delay (48 hours of roscovitine treatment) possessed just ∼2000 mutations with no evident UV-signature. Control cells that were treated by roscovitine but not exposed to UV irradiation have a highly similar spectrum of mutations and only ∼400 fewer mutations. Therefore, replication delay decreased UV-induced mutation load by more than 30 fold. This observation provides a strong support for the error-prone replication bypass of bulky lesions being the major source of mutations, at least in our experimental system. Interestingly, it also suggests that mutations in melanoma are primarily accumulating in dividing cells.

**Figure 4.**
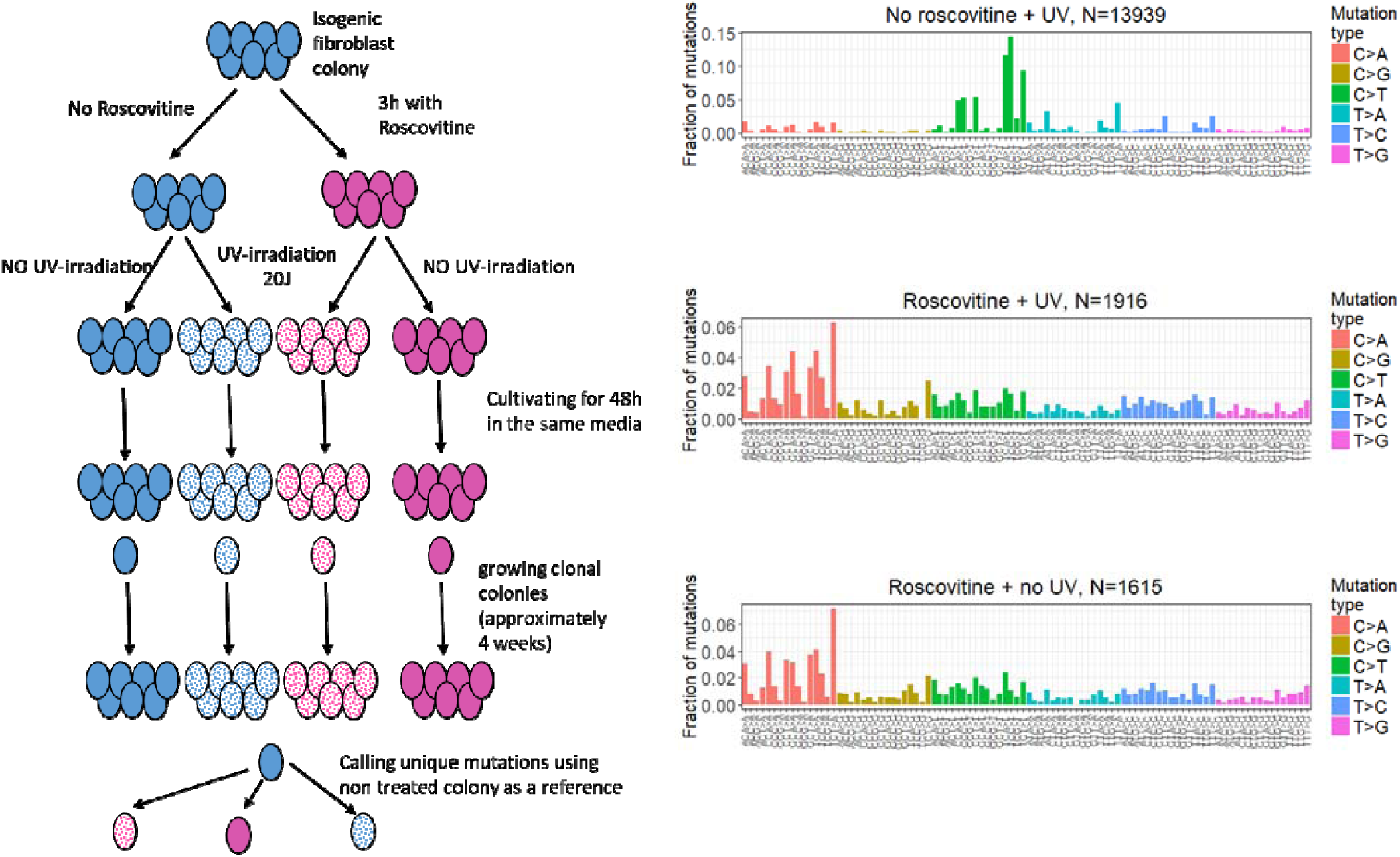
Replication delay dramatically decreases rate of UV-induced mutations. A schematic representation of the experiment is shown in the left panel: clonal colonies of fibroblast cells shown in pink were treated with roscovitine for 3 hours in advance of the UV-irradiation. Colonies shown in blue were not treated by roscovitine. Half of the colonies were irradiated with UV (20J) (dotted), and the other half were used as control. Randomly chosen cells from each colony were used to start new genetically homogeneous colonies. Numbers and spectra of mutations in these resulting colonies identified by whole genome sequencing are shown in the right panel.

R-asymmetry not related to error-prone bypass was previously detected in several cancers and in experimental systems. It was attributed to differences in fidelity between Polymerase ε and Polymerase δ^1,31–33^ or differential efficiency of mismatch repair between leading and lagging strands^31–34^. APOBEC deaminates cytosines on the lagging strand^31,33,35,36^ and misincorporation of oxo-guanin in esophageal cancer is highly asymmetric^37^. Clusters in blood cancers attributed to low-fidelity polymerase η provide another source of R-asymmetry^38^. However, these processes neither match patters observed for human germline mutations nor explain the strong association between R- and T-asymmetries and experimental data on UV-irradiated cells. It is of interest that R-asymmetry for CpG>TpG mutations in the human germline is stronger than expected for the corresponding level of T-asymmetry (Supplementary Figure 6 and Supplementary Note 1). A mechanism alternative to error-prone bypass may be responsible for R-asymmetry of these mutations that are not caused by bulky lesions.

One possible alternative explanation for the similarity between R-asymmetry and T-asymmetry in the human germline involves the exposure of DNA to a single-stranded conformation (ssDNA): the lagging strand stays in the single-stranded state during replication for a longer period, while the non-transcribed strand may occasionally adopt the single-stranded state because of R-loop formation between the transcribed strand and RNA^39,40^. We have tested the effect of R-loops on T-asymmetry and found that, in the germline, asymmetry does not increase in regions prone to R-loops compared to flanking regions within the same transcript (Supplementary Figure 7a). Additional clues to the role of ssDNA may be provided by APOBEC-induced mutations because APOBEC mutations have a strong affinity for ssDNA^41,42^. Again, we do not find that R-loops substantially affect the distribution of APOBEC-induced mutations in cancers (Supplementary Figure 7b). These analyses suggest that T-asymmetry is not mediated by ssDNA, at least as seen from R-loop data measured in human embryonic carcinoma Ntera2 cells. Consequently, it is unlikely that ssDNA is the cause of the association between T-asymmetry and R-asymmetry shown in Figure 1.

Taken together, the observed mutation patterns in the germline and in cancer, XR-seq and Damage-seq data and our experiments point to differential damage bypass rather than replication infidelity as a likely source of R-asymmetry in cancer genomes and in human germline mutations. Broadly, this suggests that DNA damage substantially contributes to spontaneous mutations. Although it is currently impossible to determine the precise proportion of damage-induced mutations, T-asymmetry allows us to conservatively quantify their contribution. Assuming that DNA damage is uniform and that TC-NER is completely error-free and is the only cause of the T-asymmetry (we correct for mutagenic effect of transcription, see Methods), we compute the minimal fraction of damage-induced mutations in highly transcribed genes. Extrapolation of this estimate to the whole genome suggests that 10% of human germline mutations (10-10%, 95% confidence intervals), 52% (51-52%, 95% confidence intervals) of mutations in melanoma, 42% (41-43%, 95% confidence intervals) of mutations in lung cancer, and 26% (24-27%, 95% confidence intervals) of mutations in liver cancer are due to DNA damage rather than replication infidelity. As expected, this estimate is much higher for cancers affected by known environmental mutagens. Still, the estimated fraction of damage-induced mutations in cancers obtained by our approach is much lower than previous estimates based on mutational spectra^8^, attesting to the conservative nature of our analysis.

From the biochemical perspective, a higher conversion rate of damage due to mutations on the lagging strand is unsurprising, as replication of the leading strand is less tolerant to damage. Helicase is attached to the leading strand and is therefore more sensitive to damage on this strand^23,25^. Furthermore, damage on the leading strand blocks Polymerase ε, which may cause fork uncoupling and stalling. This, in turn, may cause fork regression with lesion repair, template switch or homologous repair^23^ – all these processes are error-free. Fork stalling may also lead to break-induced replication resulting in highly complex mutations not analyzed here. With the exception of break-induced replication, fork stalling is usually resolved by error-free mechanisms. Meanwhile, lesions on the lagging strand are unlikely to cause fork stalling and instead often only result in a short gap downstream from the lesion^23,25^. Consequently, damage on the lagging strand is rarely removed during replication and is instead simply bypassed by error-prone mechanisms (TLS) after replication. The mutagenic effect of TLS is not limited to bypassing the damage, TLS frequently introduces mutations on the opposite strand. Our results corroborate earlier findings in the yeast system, where as much as 90% of spontaneous mutations have been attributed to TLS trough DNA lesions^43,44^.

Our experimental results show that the number of damage-induced mutations reduces with the increasing timespan between introduction of DNA damage and cell division. The computational analysis suggests that mutations statistically associated with replication do not necessarily arise as a result of replication errors alone. Several earlier studies have demonstrated the dependency of the number of accumulated mutations on the number of cell divisions. This includes dependency on paternal age for germline mutations^15–17,45^, the correlation of mutation burden in tumors with age at diagnosis^13^, and properties of the molecular clock^46^. In line with theoretical models, we note that observations showing that mutation rate scales with the number of replications do not establish the mechanistic origin of mutations^20^. Instead of being responsible for generating mutations, DNA replication may simply convert pre-existing lesions, accumulated outside of S-phase, into mutations.

## Acknowledgments

We thank Sergei Mirkin, Dmitry Gordenin, Cristopher Cassa and Donate Weghorn for useful comments on the manuscript, Lionel Sanz and Frédéric Chédin for help with R-loop data and Blake Boulerice for proofreading. This study was supported by grants: 1R35GM127131, R01MH101244 and U01HG009088; N.V.A. was supported from the Russian Science Foundation grant N°16-15-10273

## Authors Contributions

V.B.S., G.A.B. and S.R.S designed the study. V.S.B performed the data analyses. V.B.S, E.E.A, N.V.A and I.A. designed and perform experiments. M.A.A. performed data preprocessing and the helped with results presentation. S.I.N. retrieved genomic data for squamous cell carcinoma. V.B.S. and S.R.S. drafted the manuscript. All authors contributed to the final version of the paper.

## Materials and methods

### Human polymorphism and cancer mutation data

To analyze mutational patterns reflected in human DNA polymorphism, we extracted SNPs with derived allele frequency <0.1% from gnomAD data^47^. Cancer somatic mutations were extracted from PCAWG dataset^48^. Cancer somatic mutations identified in XPC wt and XPC-/-skin SCC samples were downloaded from dbGap (phs000830). Samples with MUTYH deficiency where chosen according to annotation from Scarpa et al^49^.

### Experimental data on DNA damage and repair

The XR-seq dataset for cyclobutane pyrimidine dimers (CPD) reported in Adar *et al.*^29^ allowed us to estimate the amount of DNA damage actively repaired by NER following UV irradiation. To directly assess the presence of unrepaired DNA damage, we used the Damage-seq data for CPDs provided by Hu *et al.*^30^. We did not use XR-seq and Damage-seq data for pyrimidinepyrimidone (6-4) photoproducts because these lesions are repaired too quickly to permit an accurate analysis of the effect of damage bypass over successive rounds of replication.

### R-asymmetry

As described previously^35^, the “derivative” (normalized rate of change) of replication timing may serve as a predictor of the preferential replication fork direction. This approach was proposed by Chen *et al.*^5^ and has been used in recent cancer genomics studies^31,33^.

We focused on genomic regions showing a strong preference for a specific fork direction as evident from the replication timing “derivative”. For the analysis, XR-seq, and Damage-seq (Figures 1a and b, Figures 3a and b), we used a conservative threshold corresponding to 10% of genomic regions with the highest absolute values of the replication timing “derivative”. However, this threshold appeared too restrictive for cancer genome analyses because many individual tumors have insufficient numbers of mutations within the 10% of the genome, so we relaxed the threshold to 40% for these analyses. Both of these thresholds have been used in previous studies^7,31^, and the results have generally been robust with respect to the threshold chosen.

For each individual analysis, we selected the most relevant available replication timing dataset: IMR-90 for lung cancers, HepG2 for liver cancer, and NHEK for melanoma and squamous carcinoma. For germline mutations, there is no relevant cell and we decided to consider regions with replication direction conserved across tissue types requiring that all 7 tissues have the same sign of the replication timing “derivative”; and at least in half of the tissues (4 out of 7) have value of the “derivative” exceeding 40% threshold. We also used replication timing data obtained from NHEK cell line to predict the preferential fork direction in the analysis of XR-seq and Damage-seq data and our experimental dataset (matching the tissue but not the exact cell type).

For each mutation type, we calculated R-asymmetry as the ratio of mutation density on the lagging strand to the mutation density on the leading strand. Samples with fewer than 100 mutations on each strand were excluded from the analysis to reduce sampling noise.

XPC knockouts have a distinct mutational spectrum that is dominated by Tp<u>Cp</u>T>TpTpT mutations (Supplementary Figure 8) and we restrict our test to this mutation type. Supplementary Figure 4 focuses on the magnitude of the effect in each tumor rather than on the presence of the effect. We therefore excluded samples with fewer than 500 mutations on each strand. The relaxation of the threshold to 100 mutations does not change the conclusions (data not shown).

In order to exclude the impact of T-asymmetry on the R-asymmetry estimation, we restricted the analysis of R-asymmetry to intergenic regions.

### T-asymmetry

For each mutation type, we calculated T-asymmetry as a ratio of mutation density on the transcribed strand to mutation density on the non-transcribed strand. Gene annotations and transcription direction were determined according to the knownGene track of the UCSC genome browser. Tumors with T-asymmetry >1.2 (for any of the six major mutation classes) were considered to have high level of T-asymmetry. Even with this lenient criterion, only four cancer types (melanoma, LUAD, LUSC, and liver cancer) had more than 20 tumor samples in this category. To order the genes by their expression levels, we selected the most relevant tissues from Gtex^50^: testis for SNPs from gnomAD, sun-exposed skin for melanoma, liver for liver cancer, and lung for lung cancers.

### Exclusion of replica B2 at 48h from Damage-seq

T-asymmetry and the difference between genic and non-genic regions are the main results of the Damage-seq experiments^30^ that support the utility of the data for the genome-wide analysis of bulky DNA damage and repair by the NER system. Thus, for quality control of the Damage-seq data, we calculated T-asymmetry and the ratio of reads in intergenic and genic regions separately for all replicas. T-asymmetry and the ratio of reads in intergenic and genic regions were normalized using the corresponding values for naked DNA. We found that the replicates were generally concordant at each time point with the exception of the 48h point, where we found substantial T-asymmetry and prevalence of mutations in intergenic regions in replica A but essentially no signal in replica B2 (Supplementary Figure 9). At other time points, we observed a clear, time-dependent increase in T-asymmetry and decrease in the fraction of damages in genic regions, as expected. Based on these observations, we argue that the absence of the signal in replica B2 at 48h is an artifact. Therefore, this data point was excluded. As shown in Supplementary Figure 9c, this replica is also a clear outlier in the analysis of R-asymmetry.

### Estimate of the proportion of mutations arising due to DNA damage in human cancers and the germline

To conservatively estimate the proportion of damage-induced mutations, we capitalized on the statistical signal of T-asymmetry that is associated with DNA damage. The T-asymmetry introduced by co-transcriptional processes cannot be a consequence of replication infidelity. Therefore, mutations responsible for the T-asymmetry must be damage-induced. Since transcribed and non-transcribed regions can have different susceptibilities to DNA damage, we conservatively compared the levels of mutations between transcribed strand and immediately adjacent flanking sequences rather than between transcribed and non-transcribed strands:

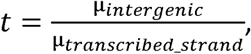

where μ_*transeribed_strand*_ is the mutation density on the transcribed strand and μ_*intergenie*_ is the mutation density in flanking intergenic regions.

To estimate *t*, we used the 10% of genes with the highest expression levels. We conservatively assumed that all damage on transcribed strands is efficiently repaired. Thus, the fraction of damage-induced mutations in transcribed regions and in intergenic regions is expressed as:

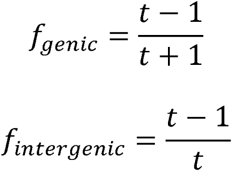

If a denotes the fraction of mutations in genic regions, and b is the fraction of mutations in intergenic regions, the fraction of damage-induced mutations for the whole genome (f_*genome*_) is expressed as:

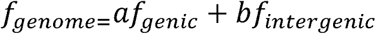

The conservative nature of this estimate is evident in the cancer data. Although nearly all mutations in melanoma are caused by UV irradiation, our estimate attributes only 50% of mutations to DNA damage (Supplementary table 3).

Confidence intervals have been obtained by sampling mutations with replacement 200 times.

### R-loops

We used data on strand-specific R-loops from Sanz *et al.* ^39^. Most R-loops were on the template strand, and we considered only such R-loops. For control regions, we used intronic regions within the same gene that were 500 nucleotides apart from the R-loop peak and 500 nucleotides long.

### CpG islands

Annotation of CpG islands was downloaded from the UCSC genome browser (cpgIslandExt).

### Experimental procedures

Human fibroblast cells from skin (GM00637) were purchased from the National Institute of General Medical Sciences Human Genetic Cell Repository (Coriell Institute). They were maintained with Minimum Essential Medium (M5650, Sigma Aldrich) supplemented with 10% fetal bovine serum (10270-106, Gibco) and 2mM L-Glutamine at 37°C in the 5% CO_2_. First, we generated genetically homogenous colonies via two successive passages starting from a single cell.

Cells were irradiated with a lamp (112537, Merck) emitting 254 nm UV light (2 J/(m2*sec)) during 10 seconds, resulting in 20 J/m2 irradiation. For a subset of colonies, we added 30µM roscovitine (R7772, Sigma Aldrich). For cells to be UV irradiated, roscovitine was added 3 hours prior to the UV treatment.

After 48 hours of incubation without changing the medium we split cells with low density in order to select individual colonies and subsequently cultivate them to achieve 1*10^6^ cells (approximately for 4 weeks). DNA was isolated with PureLink Genomic DNA Mini Kit (K182000, ThermoFisher) and then sequenced by Macrogen Inc on Illumina’s HiSeq X Ten with the average coverage of 30X. Overall, we produced six colonies including two treated with roscovitine and UV-irradiation; two UV-irradiated, but with no roscovitine in the medium; one colony incubated with roscovitine, but not irradiated; and one control colony that was not treated (Figure 4).

To quantify the change in proliferate rate after treatment with UV-light and/or roscovitine cells on coverslips were incubated with 5 µg/ml EdU for 24 hours. Then for each condition we made 15 measurements (5 different regions on 3 coverslips; at average 20 cells per region) of the fraction of cells that incorporated EdU. Cells were stained with the EdU detection kit (Click-iT EdU Imaging kit C10337, Thermo Fisher) to count divided cells and stained with Hoechst to count the total number of cells.

In each sample, we measured the proliferation rate via EdU incorporation during the first 24 hours (adding EdU 5 minutes after UV-irradiation and staining cells after 24 hours) and second 24 hours (adding EdU after 24 hours and staining the cells after 48 hours). Examples of the EdU staining are shown in Supplementary Figure 10 and Supplementary Figure 11. Adding roscovitine to the medium decreases proliferation rate by 2-3 fold compared to the control population (Supplementary Figure 12). UV-irradiation itself decreased the proliferation rate by 5 fold, followed by a substantial recovery on day 2. Combination of the UV-irradiation and roscovitine almost completely halted cell proliferation both on days one and two. Moreover, we observed that during the colony selection, cells treated with roscovitine grew slower than non-treated cells.

### Mutation calling

To obtain the set of mutations from sequenced reads, we performed following steps: first we trimmed reads with TrimGalore-0.4.5 in paired mode, then we mapped reads with bwa-0.7.12 according GATK best practice, then we call mutations from bam files with MuTect2 using the control colony (no roscovitine treatment or UV-irradiation) as “normal” and other colonies (treated with roscovitine, UV or both) colonies as “tumor”. Finally, we filtered out all the mutations observed in more than one colony. Mutation spectra for all replicates are shown on Supplementary Figure 13.

